# A three-dimensional staging system of mouse endometrial gland morphogenesis

**DOI:** 10.1101/245142

**Authors:** Zer Vue, Gabriel Gonzalez, C. Allison Stewart, Shyamin Mehra, Richard R. Behringer

## Abstract

Endometrial or uterine glands secrete substances essential for uterine receptivity to the embryo, implantation, conceptus survival, development, and growth. Adenogenesis is the process of gland formation within the stroma of the uterus that occurs after birth. In the mouse, uterine gland formation initiates at postnatal day (P) 5. Subsequently, the developing uterine glands invade into the adjacent stroma. Mouse uterine gland morphology is poorly understood because it is based on two-dimensional (2D) histological observations. To more fully describe uterine gland morphogenesis, we generated three-dimensional (3D) models of postnatal uterine glands from P0 to P21, using light sheet microscopy. At birth (P0), there were no glands. At P8, we found bud- and teardrop-shaped epithelial invaginations. By P11, the forming glands were elongated epithelial tubes. By P21, the elongated tubes had a sinuous morphology. These morphologies are homogeneously distributed along the anterior-posterior axis of the uterus. To facilitate uterine gland analyses, we propose a novel 3D staging system of uterine gland morphology during postnatal development in the mouse. We define 6 stages: Stage 0: Aglandular, Stage 1: Bud, Stage 2: Teardrop, Stage 3: Elongated, Stage 4: Sinuous, and Stage 5: Primary Branches. This staging system provides a standardized key to assess and quantify uterine gland morphology that can be used for studies of uterine gland development and pathology. In addition, our studies suggest that gland formation initiation occurs during P8 and P11. However, between P11 and P21 gland formation initiation stops and all glands elongate and become sinuous.

## Introduction

Endometrial or uterine glands are branched, tubular, epithelial structures located within the stroma of the uterus and are essential for fertility in mammals (Arora et al., 2016; Filant and Spencer, 2014). Uterine glands secrete factors that are required for uterine receptivity to the embryo, implantation, survival, development, and growth (Gray et al., 2001a). Animal models that lack uterine glands are infertile because they are unable to produce histotroph. Infertility occurs in these models due to defects in embryo implantation and early pregnancy loss (Cooke et al., 2012; Dunlap et al., 2011; Filant et al., 2012; Gray et al., 2001a; Gray et al., 2001b; Gray et al., 2002; Spencer and Gray, 2006; Spencer et al., 2005).

Uterine gland development is a unique developmental process that occurs after birth. In the mouse, uterine adenogenesis (gland formation) initiates at postnatal day (P) 5 (Cooke et al., 2012). At P5, there are invaginations of the luminal epithelium of the uterus into the adjacent mesenchyme. Subsequently, developing endometrial glands extend into the surrounding stroma (Spencer et al., 2005). By P15, the histoarchitecture of the uterus has developed into its mature form with endometrial and myometrial layers (Hu et al., 2004).

Rodent uterine glands are considered simple tubular structures that are not tightly coiled or extensively branched as in other mammalian species. In neonatal rodents, the initial formation of uterine glands is ovary and steroid independent (Cooke et al., 2012; Filant et al., 2012; Spencer et al., 2006; Stewart et al., 2011). However, the post-pubertal period of uterine gland development is hormone dependent (Stewart et al., 2011; Wetendorf and DeMayo, 2012; Wetendorf and DeMayo, 2014). During homeostasis, the epithelial and stromal compartments of the mouse uterus remain separate. However, after parturition, the uterine glands can regenerate through a stromal to epithelial transition (Huang et al., 2012; Patterson et al., 2013).

Although uterine glands have been studied for decades, there is very limited information on their three-dimensional (3D) structure and no volumetric information on their 3D structure during development (Arora et al., 2016; Cooke et al., 2012). Current models for the 3D structure of uterine glands rely on interpretations of 2D histology (Gray et al., 2001a; Hu et al., 2004). Three-dimensional structures can be determined by reconstructions of serial histological sections of uterine glands but it is time-consuming, labor-intensive, and until recently resulted in relatively crude structures (Hondo et al., 2007; Nunobiki et al., 2001). Recent advances in the 3D imaging of biological specimens and subsequent volumetric analyses have enabled the visualization and quantification of whole embryos, including developing organs and tissues (Belle et al., 2017; Hsu et al., 2016; Short et al., 2014).

We have developed a new method to image and render the 3D structure of developing uterine glands in the mouse. This was achieved using whole mount immunofluorescence for an epithelial marker, tissue clearing, and light sheet microscopy. Light sheet or single selective-plane illumination has a distinct advantage over other fluorescent microscope systems due to multi-angle image acquisition and 3D reconstruction of optical sections. In addition, the ability to rotate samples reduces limitations of depth penetration due to light scatter. Analysis of the 3D epithelial structures during adenogenesis revealed the dynamics of glandular morphogenesis. We find that there is an initial period of gland formation initiation that is followed by a period of gland differentiation and morphogenesis. Using our findings, we have created a novel 3D staging system for mouse uterine glands. Our uterine gland staging system should be useful in the future to quantify uterine gland mutant phenotypes and pathologies.

## Results

### 3D models of uterine glands within the developing postnatal mouse uterus

Uterine glands are believed to be derived from the luminal epithelium and remain connected with the lumen once formed (Gray et al., 2001a). However, in 2D histological tissue sections, uterine gland tissue can be seen as discrete round or oval epithelial units within the stromal compartment, not continuous with the luminal epithelium (Filant et al., 2012). To determine the morphology of mouse uterine glands in three dimensions, we performed whole mount immunofluorescence on individual postnatal uterine horns with an epithelial-specific antibody TROMA-1 (cytokeratin 8/18), that was followed by tissue clearing and light sheet microscopy. This approach allowed us to visualize the 3D morphologies of the uterine epithelium at postnatal ages during adenogenesis. These postnatal ages included postnatal day (P) 0 (P0), P8, P11, and P21, based on insights from previous 2D histological studies (Fig. 1).

**Fig. 1.**
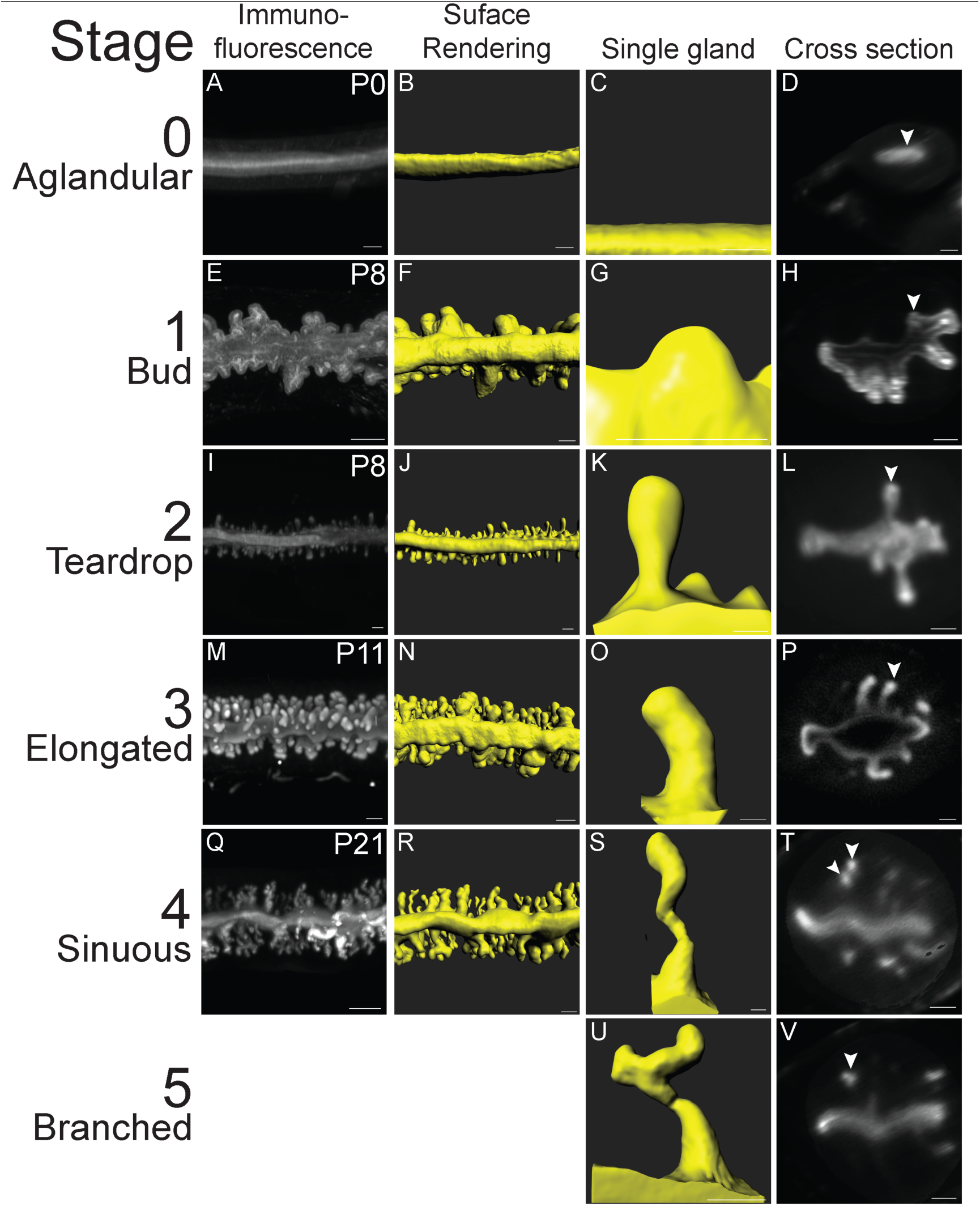
3D image analysis of the postnatal mouse uterus. Fluorescent light sheet images and 3D reconstructions of the postnatal mouse uterus and individual uterine glands. (A, E, I, M, Q) TROMA-1 stained immunofluorescent images of the uterine epithelium. Mesometrial view. (B, F, J, N, R) Pseudo-colored surface renderings of the uteri corresponding to the first column. Mesometrial view. (C, G, K, O, S, U) Pseudo-colored surface renderings of single uterine glands. (D, H, L, P, T, V) Optical cross-sections of fluorescent light sheet images of TROMA-1 stained uteri. Arrowheads correspond to the gland in the third column. Anti-mesometrial to the right. (A-D) P0, (E-L) P8, (M-P) P11, and (Q-V) P21. Scale bars: 100 μm.

The TROMA-1 antibody labels luminal and glandular epithelial cells in the uterus (Brulet et al., 1980). Volumetric fluorescent images were acquired and rendered as 3D reconstructions (Fig. 1A, E, I, M, Q). These 3D reconstructions revealed the gross structure of the uterine epithelium at these postnatal stages. At this level of analysis, at birth (P0), adenogenesis has not initiated (Fig. 1A). There was only a smooth luminal epithelium (Fig. 1B-D). However, at P8, P11, and P21 we observed forming uterine glands (Fig. 1E, I, M, Q). Thus, adenogenesis commences between P0 and P8. Surface renderings of the 3D reconstructions of the fluorescent images provide a simpler view of the uterine epithelial compartment and more specifically the structures of individual uterine glands (**Fig. B, F, J, N, R**).

### Distinct uterine gland morphologies are associated with postnatal age

To visualize the glandular epithelium and individual uterine gland morphologies without the luminal epithelium fluorescent signal, we used the FilamentsTracer software to segment the glandular epithelium from the luminal epithelium. We manually marked each gland as a “dendrite” in each z-slice in the z-stack of each volumetric image. The region of the gland nearest the lumen and luminal epithelium was marked as the base, while the region of the gland nearest the myometrium was marked as the end. Once each gland was marked, we generated volumetric surface renderings of the forming glands, based on the fluorescent signal. This allowed us to eliminate the luminal epithelial fluorescent signals. The computationally-isolated uterine glands could then be assessed for quantitative measurements.

The 3D surface rendering of the uterine glands revealed a set of distinct morphologies associated with specific postnatal ages. At P0 (n=6 uterine horns), the mouse uterus contained no glands (Fig. 1A-D, **Movie 1**), verifying that uterine gland morphogenesis is a postnatal developmental process. At P8 (n=5), there were rounded extensions of the luminal epithelium into the stroma that we refer to as “buds” (Fig. 1E-H, **Movie 2**). Bud stage uterine glands have a diameter greater than their length. At P8, there were also forming uterine glands with a teardrop-shaped morphology (Fig 1I-L, **Movie 3**). Teardrop stage uterine glands have a constricted region adjacent to the luminal epithelium and a more rounded appearance extending toward the forming myometrium. The volume (epithelium and lumen of the gland) of P8 bud-stage uterine glands (n=109) was 0.3 ± 0.2 × 10^5^ um^3^, whereas the volume of P8 teardrop-stage uterine glands (n=152) was 1.4 ± 0.6 × 10^5^ um^3^ (Fig. 2). Thus, at P8, adenogenesis has initiated and there are two distinct uterine gland morphologies.

**Fig. 2.**
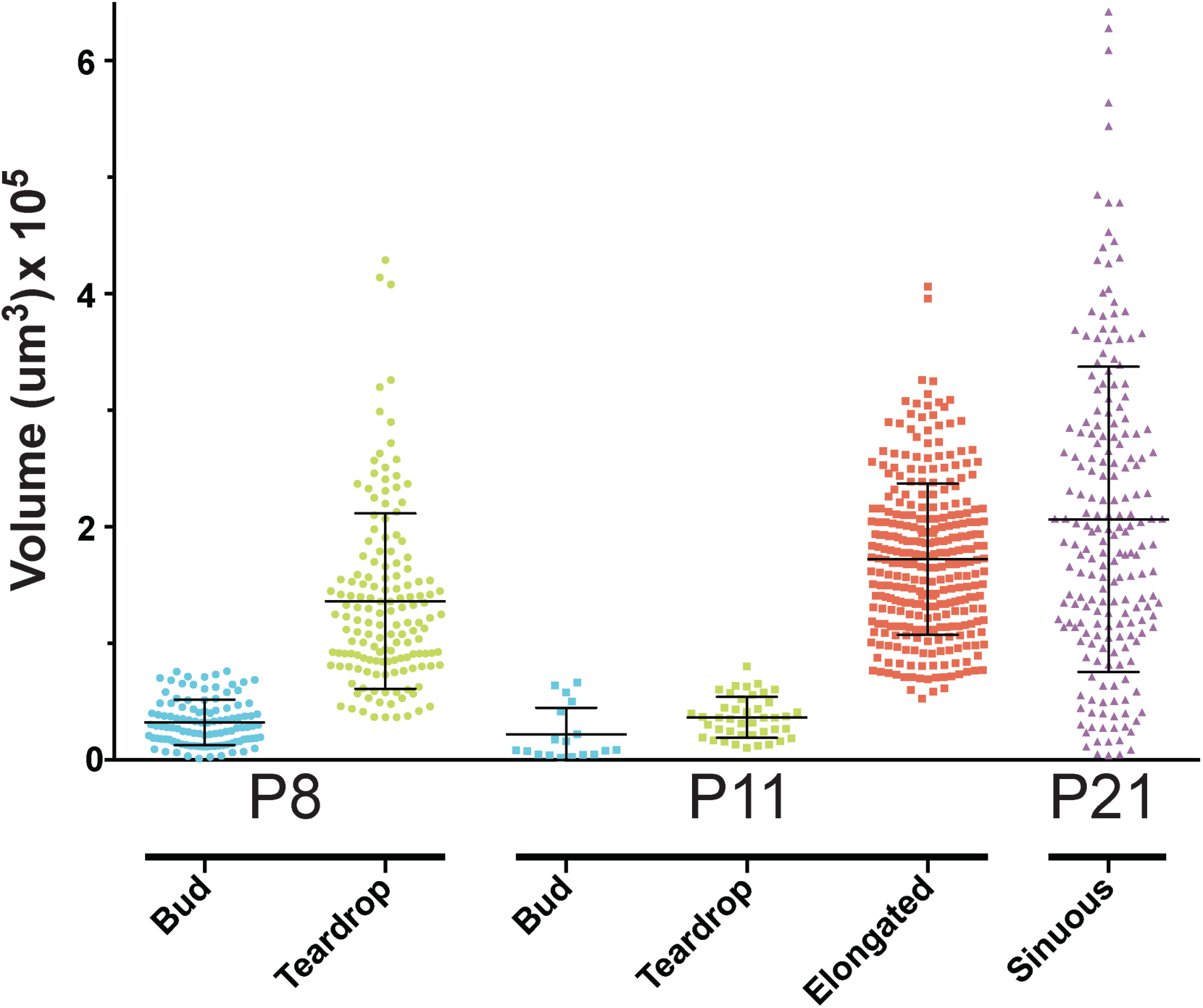
Uterine gland volume during postnatal development. Glands were measured with Imaris FilamentsTracer to quantify the volume of various gland types (bud, teardrop, elongated or sinuous) at P8, P11 and P21. Each dot, square or triangle represents the volume of an individual gland. Error bars indicate average volume ±1 standard deviation.

At P11 (n=5), bud- and teardrop-shaped glands were present (Fig. 1M-P). In addition, there were elongated glands that extended towards the inner circular myometrial layer (Fig. 1M-**P**, **Movie 4**). Elongated glands are defined as being longer than bud- and teardrop-shaped uterine glands without the luminal constriction associated with teardrop-shaped glands. The average volume of bud-stage glands (n=18) at P11 is 0.2 ± 0.5 × 10^5^ um^3^, teardrop-shaped glands at P11 (n=42) is 0.4 ± 0.2 × 10^5^ um^3^, and elongated glands at P11 (n=298) is 1.7 ± 0.4 × 10^5^ um^3^ (Fig. 2).

At P21 (n=3), when the uterine histoarchitecture is similar to that of the adult uterus, the glands are sinuous and coiled, extending from the uterine lumen to the inner circular myometrium (Fig. 1Q-T, **Movie 5**). An optical section of a sinuous gland shows that there can be multiple discrete round/oval epithelial units from a single gland in a z-stack slice (Fig. 1T). Rarely (0.5%), some uterine glands are branched (Fig. 1U-V). At P21, only sinuous uterine glands and rare branched glands were present. No bud-, teardrop-shaped or elongated uterine glands were found. The average volume of P21 glands (n=204) is 2.0 ± 0.9 x 10^5^ um^3^ (Fig. 2). P21 glands had the highest variation in terms of volume compared with the other uterine gland morphologies (Fig. 2). Indeed, sinuous glands with larger and smaller volumes can be observed in the surface rendered images (**Movie 5**).

In addition to morphology, it is possible to segregate the different uterine gland structures by volume. At P8, bud- and teardrop-stage uterine gland volumes have minimal overlap (Fig. 2). However, at P11, there is considerable overlap in volume between the bud and teardrop uterine gland morphologies and therefore must be sorted by their structure (Fig. 2). Whereas elongated glands had a wide range of volumes, there was very little overlap with bud- or teardrop-shaped gland volumes (Fig. 2).

### Uterine gland morphologies are homogeneously distributed along the anterior-posterior axis

To visualize uterine gland morphology distribution along the anterior-posterior axis, individual uterine horns were isolated at P8 and P11 for volumetric imaging. Multiple images were collected to recover the majority of the uterine horn. At P8, we were able to acquire images from a central region of the uterine horn encompassing ~80% of the organ. At P11, we were able to acquire images from a central region of the uterine horn encompassing ~60% of the organ.

At P8, the distribution of uterine glands was examined by morphology (bud versus teardrop) and volume along the anterior-posterior axis (Fig. 3A). We found that bud-shaped glands and teardrop-shaped glands were homogeneously distributed along the anterior-posterior axis of the uterine horn. We found that the volume of buds was below 1 × 10^5^ μm^3^, while the majority of the volumes of teardrop-shaped glands were above this value. However, there was some overlap in volume between bud- and teardrop-shaped glands. In addition, teardrop-shaped glands have a higher variation in volume compared to bud-shaped glands.

**Fig. 3.**
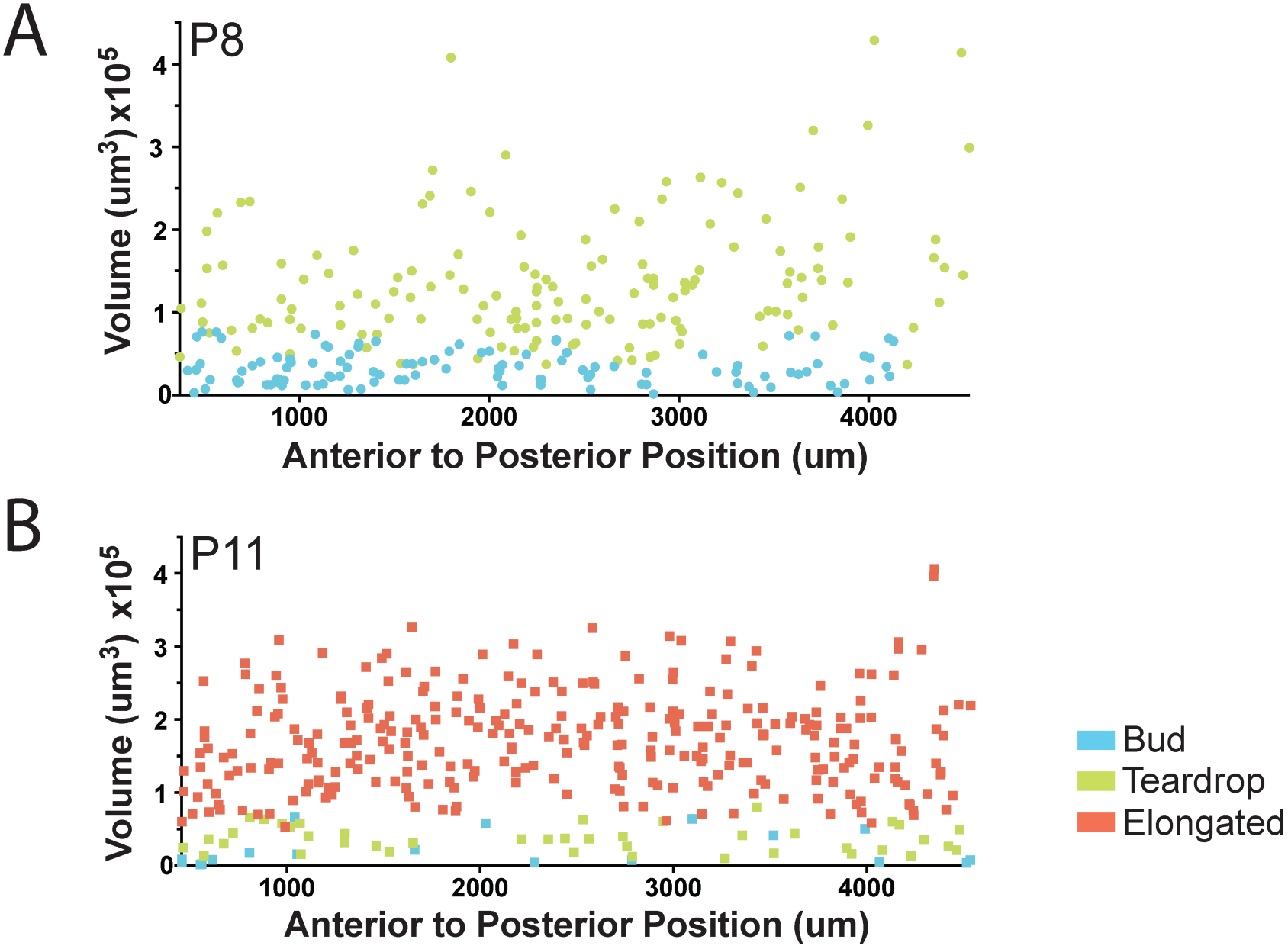
Uterine gland distribution in the postnatal uterus. Uterine gland morphology and volume along the anterior-posterior axis of the uterine horn. (A) P8, buds (blue circles), teardrops (green circles) (B) P11, buds, (blue squares), teardrops (green squares), elongated (red squares).

We also examined the distribution of uterine glands at P11 by morphology (bud versus teardrop versus elongated) and volume along the anterior-posterior axis of the uterine horn (Fig. 3B). Buds and teardrops are present at P11. However, there are substantially fewer bud- and teardrop-shaped glands compared to elongated glands. We found that all three types of gland structures: buds, teardrops, and elongated glands were distributed homogeneously along the anterior-posterior axis of the uterine horn. Thus, there does not appear to be a spatially-restricted pattern of uterine gland morphologies along the anterior-posterior axis of the uterine horn at P8 and P11.

### Identification of the uterine rail, a novel structure in the mouse uterine horn

3D imaging of the uterine horns also allowed us to identify gross morphological features not previously appreciated by 2D histology (Fig. 4). We found a structure in the aglandular region located along the mesometrial pole of each uterine horn. This region was not present at P0 but was found at P8, P11, and P21 (Fig. 4A-H, Movies 1–4) but. The 3D structure of this uterine feature appears similar in morphology to a railroad track (Fig. 4D, F, H). Thus, we have termed this structure the “uterine rail”.

**Fig. 4.**
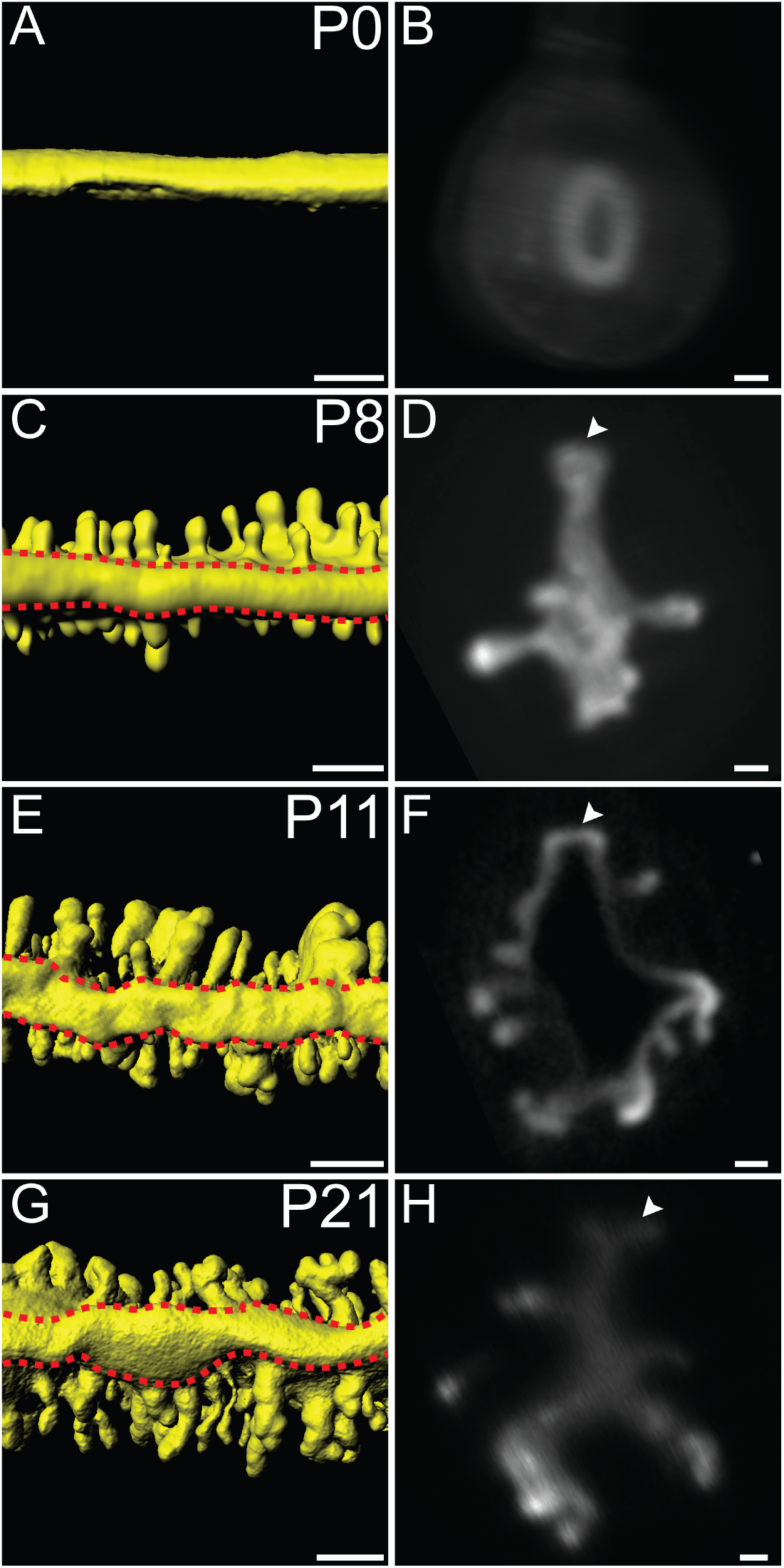
Identification of the uterine rail. 3D reconstructions of the TROMA-1 stained postnatal mouse uterine epithelium. Mesometrial views. (A, C, E, G) Red dashed lines outlines the uterine rail. (C, E, G) Optical cross-sections of fluorescent light sheet images of TROMA-1 stained uteri. Arrowheads, uterine rail. Mesometrial, top. (A, B) P0, (C, D) P8, (E, F) P11, and (G,H) P21. Scale bars for (A, C, E, G) = 150 μm. Scale bars for (B, D, F, H) = 50 μm.

When visualizing the epithelial compartments of the uterine horns at various postnatal stages, we also observed morphological differences in the shape of the uterine lumen. At P8 and P11, the uterine lumen has a relatively straight morphology when viewed from the mesometrial pole (Fig. 1F, J, **N, Movie 2, 3). However, at P21, the uterine lumen has a zig-zag morphology (Fig. 1R, Movie 4**).

## Discussion

We have generated 3D reconstructions of the epithelial compartment of the developing postnatal mouse uterus. This was accomplished using whole mount immunofluorescence, tissue clearing, light sheet microscopy and computational processing. Our study shows for the first time the 3D morphology of individual developing endometrial glands and their distribution within the intact uterus. Previously, 2D histological sectioning has been used to infer 3D uterine gland morphologies during adenogenesis (Branham et al., 1985; Hu et al., 2004). In some cases, subsequent 3D reconstructions were employed to examine uterine gland morphology (Hondo et al., 2007; Nunobiki et al., 2001). However, deconstruction of the tissue by histological sectioning is labor intensive and at the time 3D reconstruction abilities were relatively crude. Optical sections of a whole mount organ do not require tissue deconstruction. In addition, the subsequent 3D rendering of these optical sections provides an opportunity to generate computational sections of the organ at any desired plane. These advantages have provided views of uterine gland development in their natural state within the organ intact. Although the FilamentsTracer software was created for the automated detection of neuronal structures, we successfully used it to segment developing uterine glands (Swanger et al., 2011). It is likely that this software could be used to segment non-neuronal structures in other developing tissues. Recently, 3D reconstructions of adult mouse endometrial glands have been achieved, showing their complex branched morphologies (Arora et al., 2016).

### Glandular development in the mouse uterus

At birth (P0), the uterus consists of a simple epithelium surrounded by undifferentiated mesenchyme that lacks endometrial glands (Hu et al., 2004). Our 3D imaging verifies these observations. By P5, the mesenchyme has developed into three separate layers: the radially oriented endometrial stroma, the inner circular myometrial layer, and the prospective outer longitudinal myometrium. Fluorescent images of uterine epithelium enzymatically and mechanically isolated from the stroma and myometrium at P5 show numerous epithelial buds (Cooke et al., 2012). However, these images were not rendered in 3D. In addition, there were likely morphological distortions of these structures during epithelial isolation. Our results at P8 are consistent with these findings but also provide 3D structural information of individual glands and their distribution in the uterine horn. Indeed, we could distinguish bud- and teardrop-shaped morphologies. Histological studies of mouse uteri at P12 show the presence of glandular epithelium but it is not possible to distinguish individual gland morphologies (Hu et al., 2004). At P11, our 3D analysis identified long, straight, tubular glands as well as bud- and teardrop-shaped glands. Fluorescent images of uterine epithelium enzymatically and mechanically isolated from the stroma and myometrium at P20 show a network of glandular tissue but individual gland morphologies are not distinct (Cooke et al., 2012). Our 3D imaging at P21 showed the sinuous and coiled nature of the uterine glands and the absence of the bud, teardrop and elongated glands.

Our data are consistent with a model for uterine gland formation (Fig. 5). Uterine gland formation initiates by forming an epithelial bud into the surrounding stroma. The bud then develops into a teardrop-shaped structure. The teardrop-shaped gland will extend further into the stroma to form an elongated gland. As elongated glands continue to lengthen, they will become sinuous and coiled. Subsequently, the uterine gland will branch. Time-lapse imaging could be used to test this model. However, in vitro culture systems for the postnatal mouse uterus are limited for volumetric imaging over many days (Newbold et al., 1994).

**Fig. 5.**
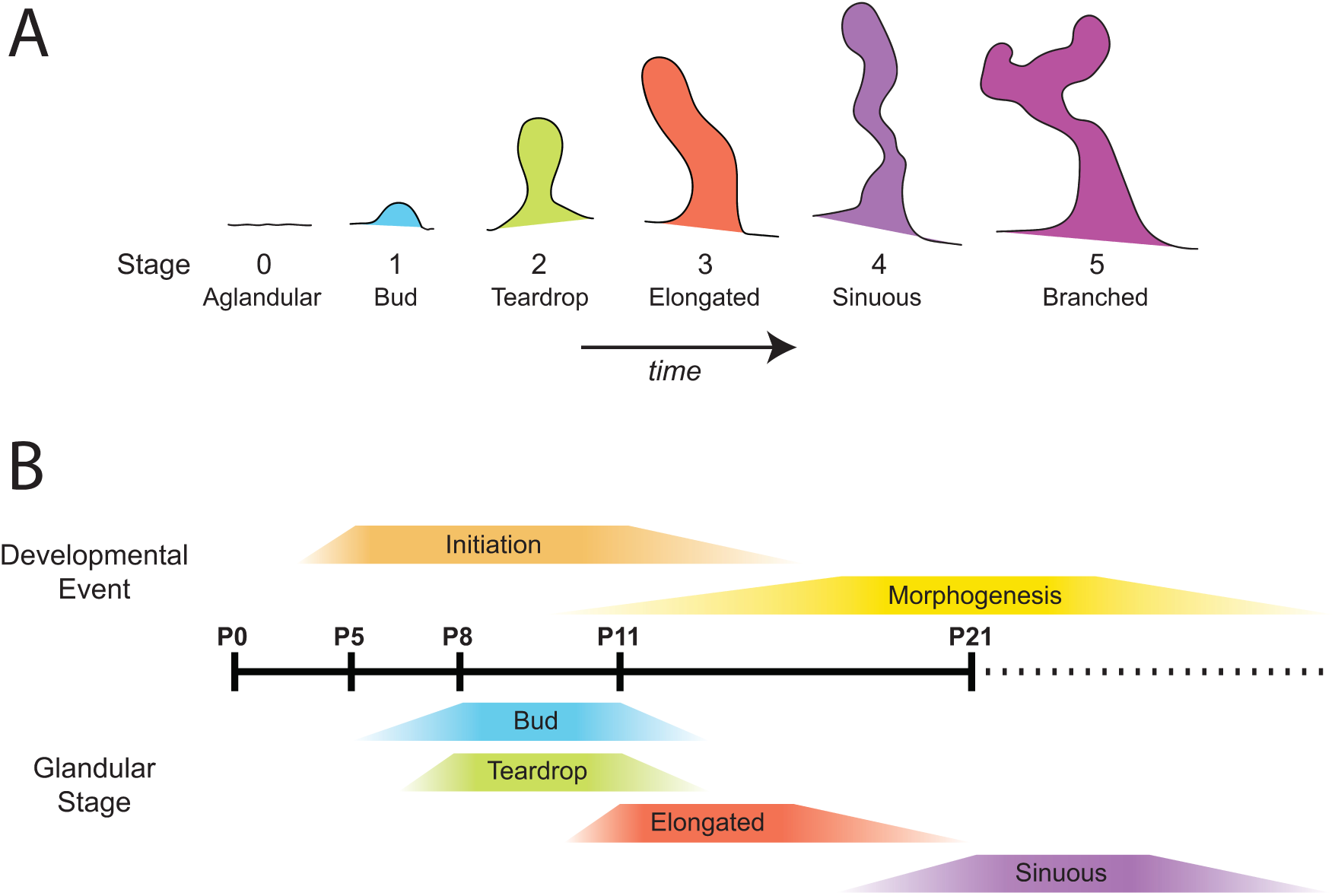
Uterine gland staging system and development. (A) Uterine gland staging system. Stage 0 = Aglandular, no glands are present. Stage 1 = Bud. Stage 2 = Teardrop. Stage 3 = Elongated. Stage 4 = Sinuous. Stage 5 = Branched. (B) Correlation of gland formation initiation and morphogenesis with postnatal age.

Previous studies have shown the presence of forming uterine glands at P5 (Cooke et al., 2012). We have also performed 3D imaging of P5 uteri and observe bud-shaped glands (Vue and Behringer, unpublished observations). Our 3D studies identified bud-shaped glands at P8 and P11 but not at P21. This suggests that gland initiation occurs during the period from P5 to P11. Between P11 and P21, gland initiation stops and switches to terminal morphogenesis because, by P21, there are no bud, teardrop or elongated glands. Interestingly, there were sinuous glands with large volumes and small volumes. The small volume glands are relatively thin compared to the large volume glands. It is possible that there may have been some glands at the bud and teardrop or early elongated stages that were present when a terminal differentiation signal was present, leading to a sinuous morphology but small volume.

We were able to 3D image individual forming uterine glands and also their distribution within the uterine horn. We found that bud, teardrop, elongated and sinuous gland morphologies were distributed homogeneously along the anterior-posterior axis of the uterine horn at P8 and P11. This suggests that there is no anterior-posterior pattern for uterine gland development. This implies that mechanisms for uterine gland formation are distributed homogeneously from anterior to posterior. However, we did observe a novel structure dorsal structure located on the mesometrial side of the uterine horn we call the uterine rail. This region is known to be free of uterine glands but to our knowledge, the uterine rail structure has not been described previously. This may be because uterine sections are typically placed on slides in any orientation. We suggest that uterine horn sections be placed in a consistent manner with the mesometrial side placed up. However, this highlights the benefits of 3D renderings of volumetric imaging.

In our study, we employed outbred Swiss Webster mice. The present study indicates that uterine adenogenesis in Swiss Webster mice generally occurs during the same time period as previously described for BALB/c and C57BL6 mice (Brody and Cunha, 1989; Cooke et al., 2012). However, there may be timing differences in adenogenesis between outbred and various inbred mouse strains.

### A novel 3D staging system of uterine gland development in the mouse

Our description of the 3D structures of individual uterine glands provides a framework for examining adenogenesis during normal and pathological situations. Thus, we propose a novel 3D staging system of mouse uterine glands, dividing gland morphologies into six stages (Fig. 5A).

**Stage 0** – Aglandular Stage. No glands or forming glands are present. Stage 0 occurs at P0. **Stage 1** – Bud Stage. Bud-shaped glands are relatively shallow epithelial invaginations into the adjacent stroma. Buds are defined as having a larger diameter compared to its length. They are found at P8 and P11. **Stage 2** – Teardrop Stage. Teardrop-shaped glands are similar to buds but their length is longer than their diameter. In addition, there is a narrowing of the gland closest to uterine lumen. They are found at P8 and P11. **Stage 3** – Elongated Stage. Elongated glands have a longer length than diameter but do not have the teardrop narrowing at the uterine lumen. They are present at P11. **Stage 4** – Sinuous Stage. Sinuous-shaped glands are elongated but also curved and coiled. They are present at P21. **Stage 5** – Branched Stage. Stage 1–4 glands do not have branches. Stage 5 glands are branched. These glands are present as early as P21 and are also present in adults (Arora et al., 2016).

Attempts to quantify uterine gland defects in mouse mutants have been challenging because in 2D histological sections it is not clear where one gland begins and another ends. Indeed, we have used numbers of gland cross sections as a surrogate to quantify glandular content in the uterus (Gonzalez et al., 2016; Stewart et al., 2011; Stewart et al., 2013). 3D imaging of developing uterine glands combined with our proposed staging system should lead to more precise assessments of endometrial pathologies.

## Materials and Methods

### Mice

Swiss Webster outbred mice (Taconic Biosciences) were used in this study. All mice were maintained in compliance with the Public Health Service Policy on Humane Care and Use of Laboratory Animals, the U. S. Department of Health and Human Services Guide for the Care and Use of Laboratory Animals, and the United States Department of Agriculture Animal Welfare Act. All protocols were approved by the University of Texas MD Anderson Cancer Center Institutional Animal Care and Use Committee.

### Immunostaining

Female reproductive tract organs were dissected and placed into 4% paraformaldehyde (PFA) on a nutator for 16 hours at room temperature. The next day, tissues were washed twice with phosphate buffered saline (PBS) for an hour, and transferred into DMSO (dimethyl sulfoxide) and methanol (1:4 ratio) and stored at -20°C overnight or longer.

For whole mount immunostaining, samples were washed in 50% methanol for an hour, then in PBS for an hour and transferred into 50 ug/mL proteinase K for 5–20 minutes. To stop the reaction, samples were placed back into PBS. To permeabilize the tissue, samples were transferred into clearing solution (200 mM boric acid, 4% sodium dodecyl sulfate, pH 8.5) on a nutator for 1 to 16 hours at room temperature. Samples were washed in PBST (0.1% Triton X-100 in PBS) overnight and then placed into blocking solution (10% normal goat serum and 1% Triton-X in PBS) overnight on a nutator at room temperature. Primary antibody TROMA-1 (Developmental Studies Hybridoma Bank, Iowa USA, 1:100) was added and incubated at room temperature for 3 days. Samples were transferred into PBS for 6 hours at room temperature, changed into blocking solution for 1 hour at room temperature, placed into secondary antibody (AlexaFluor, BD Biosciences, San Jose, CA, USA, 1:200) for 3 days at room temperature and back into PBS for 6 hours. Samples were then fixed in PFA for an hour at room temperature and washed in PBS for at least an hour. To clear the tissues, samples were washed in Sca*l*eA2 (4M urea, 10% glycerol and 0.1% Triton X-100) overnight at room temperature. Uteri were cut in half to isolate individual horns and embedded in 1% low melt agarose in a capillary tube (100 μl, inner diameter 2.5 mm, Brand GMBH, catalog number 701910). The agarose-embedded uterine horn was then transferred into a petri dish containing Sca*l*eA2 and placed on a nutator overnight.

### Image capture and post-acquisition processing

Whole-mount fluorescent images were obtained using a Zeiss Lightsheet Z.1 Light Sheet Fluorescence Microscope (LSFM) (Zeiss, Jena Germany) with 405/488/561/640 nm lasers. For all light sheet images, a 5×, NA0.6 Plan-Apochromat water immersion objective (Carl Zeiss) and dual-sided illumination were used. Laser power and exposure times varied depending on the amount of fluorescent signal and proximity of the signal to the surface of the tissue. All Z-stacks were processed through Imaris (Bitplane), including pseudo-coloring, adjusting the dynamic range of each color channel, surface rendering, background subtraction and segmentation of specific tissues. To assess the volume and position of individual glands, FilamentTracer software (Imaris) was adapted for our use. FilamentsTracer was designed to detect neurons (dendritic trees, axons, and spines), microtubules and other filament-like structures. The analysis was performed by manually demarcating the base and tip of each gland and designating each gland as a “dendrite” by using the default settings. Volume and uterine position calculations were determined using FilamentTracer software.

### Statistical Analysis

Statistical analysis was performed using the Prism 6.0 software (GraphPad). The frequency distribution of glands was performed using column analyses and column statistics for all postnatal time points. For P8 glands (bud versus teardrop), a two-tailed Student T-test, unpaired, non-parametric test was performed. For P11 (bud versus teardrop versus elongated) were analyzed using a one-way ANOVA. For statistical analyses, α=0.05 was used to determine significance.

## Acknowledgements

We thank Chih-Wei Hsu and Tegy John Vadakkan from Baylor College of Medicine Optical Imaging and Vital Microscopy Core for microscopy and assistance, Adriana Paulucci and Henry Adams from the MD Anderson Department of Genetics Microcopy Core for microscopy and post-image acquisition processing, and Kenneth Trimmer for help with Imaris software. We are grateful to Jichao Chen for initial imaging by Optical Projection Tomography. This research was supported by National Institutes of Health (NIH) grant HD030284 and the Ben F. Love Endowment to R.R.B. Z.V. was supported by NIH NIGMS R25GM56929, NIH T32-CA009299-27, and NIH HD30284. Veterinary resources were supported by NIH grant CA16672.

---

**Movie 1.** TROMA-1 immunofluorescence and surface-rendered opaque view of a P0 uterine horn.

**Movie 2.** TROMA-1 immunofluorescence and surface-rendered opaque view of a P8 uterine horn.

**Movie 3.** TROMA-1 immunofluorescence and surface-rendered opaque view of a P11 uterine horn.

**Movie 4.** TROMA-1 immunofluorescence and surface-rendered opaque view of a P21 uterine horn.

**Movie 5.** TROMA-1 immunofluorescence and surface-rendered opaque view of an individual “bud” stage uterine gland.

**Movie 6.** TROMA-1 immunofluorescence and surface rendered opaque view of an individual “teardrop” stage uterine gland.

**Movie 7.** TROMA-1 immunofluorescence and surface-rendered opaque view of an individual “elongated” stage uterine gland.

**Movie 8.** TROMA-1 immunofluorescence and surface-rendered opaque view of an individual “sinuous” stage uterine gland.

